# ACC1-Dependent De Novo Lipogenesis Sustains Hematopoietic Stem Cell Quiescence and Self-Renewal

**DOI:** 10.64898/2026.06.03.729986

**Authors:** Morgan A. Jones, Andrew DeVilbiss, Timothy A. Liang, Sho Matono, Zhiyu Zhao, Adam Ross, Daniel Cassidy, Sean J. Morrison, Qing Li

## Abstract

Certain aspects of lipid metabolism are known to regulate hematopoietic stem cell (HSC) function, including fatty acid oxidation and lipid uptake, but there is a limited understanding of the contribution of de novo fatty acid synthesis to HSC homeostasis. Here, we show that endogenous fatty acid synthesis is essential for HSC function. Conditional deletion of *Acaca*, the gene that encodes the rate-limiting enzyme for de novo fatty acid synthesis, acetyl-CoA carboxylase 1 (ACC1), in hematopoietic cells profoundly reduces HSC function, marked by a reduced ability to reconstitute irradiated mice after competitive transplantation. *ACC1* deficiency reduced quiescence, increased uptake of extracellular lipids, and increased reactive oxygen species in HSCs. The loss of HSC function is partly caused by increased fatty acid oxidation as deletion of CPT1a, which is required for long-chain fatty acid oxidation, partially rescued HSC function. A balance between fatty acid synthesis and fatty acid oxidation is thus critical for the maintenance of HSC function.

## Introduction

Hematopoietic homeostasis requires that a rare pool of hematopoietic stem cells (HSCs) balance the processes of differentiation and self-renewal to facilitate the production of hundreds of billions of blood cells every day. This process must be adaptive such that production is amplified in the setting of blood loss or infection and thus requires both cell intrinsic signals and extracellular cues to balance hematopoiesis. It is becoming increasingly evident that metabolic plasticity is an essential feature of HSC adaptive capabilities, though the precise metabolic programs and contexts in which they may be important are largely unknown.

Fatty acids (FAs) are essential for membrane building, signaling, and bioenergetics, but HSC capabilities surrounding manufacture and general utilization of FAs remain poorly characterized. Work to date has identified that under infectious stress, HSCs import extracellular FAs to generate energy through β-oxidation.^1^ In addition, leukemic stem cells also rely on extracellular fatty acids, and the uptake of fat may be central to chemoresistance in at least some leukemias.^2^ However, whether extracellular FAs are required for normal HSC function under steady state remains unclear. When cultured in serum-free culture conditions, HSCs are deprived of exogenous FAs and upregulate FA biosynthetic machinery, suggesting the importance of FAs to HSC function *ex vivo*.^3^ However, whether FA synthetic machinery is important to steady state HSC homeostasis *in vivo* has not been investigated.

On the other hand, recent studies have demonstrated that FA catabolism is essential for HSC function *in vivo*. Inhibition of β-oxidation with the drug etomoxir impacted the ability of HSCs to maintain self-renewal.^4^ While this suggests a fundamental role of β-oxidation in HSC biology, etomoxir has targets beyond carnitine palmitoyltransferase 1A (CPT1A), the gatekeeper to β-oxidation, and furthermore may be directly toxic to cells.^5,6^ Recent work in which CPT1A was directly inactivated in HSCs through mouse genetic knockout models demonstrated that HSCs became dependent on β-oxidation through CPT1A as they aged, reflective of an erosion of HSC metabolic plasticity resultant from the aging process.^7^ Intriguingly, this same study demonstrated that under the stress of high fat feeding, engagement of mitochondrial metabolism through CPT1A was detrimental to HSC function, and CPT1A loss was therefore protective, thus indicating a context dependent impact of activation of mitochondrial β-oxidation.^7^ Mitochondrial catabolism of FAs is therefore essential to HSCs, but the source of these FAs remains unknown.

FAs can either be transported into HSCs or synthesized internally, though the latter pathway, termed *de novo* lipogenesis (DNL), has not been identified as active in HSCs *in vivo*. DNL is an energetically demanding process known to be active in the liver and in adipose tissue.^8^ DNL is catalyzed by two proteins. The first rate-limiting step of DNL is the carboxylation of acetyl-CoA to form malonyl-CoA by the protein Acetyl-CoA Carboxylase 1(ACC1; encoded by *Acaca*). Malonyl-CoA is then extended through a series of condensation reactions with additional acetyl-CoA molecules to form palmitate through the function of Fatty Acid Synthase (FASN; encoded by *Fasn*). DNL activity has not been queried in steady state HSCs to date, and it is unclear the extent to which this pathway is active in somatic stem cells in general.

Here, we demonstrate that de novo lipogenesis is indispensable for HSC maintenance and self-renewal. Conditional deletion of *Acaca*, the gene encoding ACC1, results in profound HSC dysfunction characterized by loss of quiescence, impaired regenerative capacity, and elevated oxidative stress. HSC function is partially rescued by inhibition of mitochondrial FAO. Our findings identify de novo lipogenesis as a fundamental metabolic pathway that preserves HSC integrity and reveal an unexpected requirement for endogenous FA synthesis in the maintenance of stem cell quiescence and long-term regenerative potential.

## Results

### Primitive hematopoietic cells have increased unsaturated fatty acids (UFAs) compared to downstream progenitors

The quantitation of metabolites in rare cell populations, most notably the ultrarare hematopoietic stem and multipotent progenitor cell populations (HSC/MPP; CD48-LSK) has been difficult due to cell number requirements, the *ex vivo* time required to obtain adequate cell numbers, and environmental contamination limiting analysis of lower abundance metabolites. To overcome this limitation, we utilized a rigorous, clean, and rapid approach previously described.^9,10^ In comparing HSC/MPPs to more differentiated hematopoietic progenitor cell 1/2s (HPC1/2; CD48+LSK), we found that the primitive HSC/MPP compartment was enriched for unsaturated fatty acids (UFAs) (**Figure 1A**). This suggests the possibility of increased UFA uptake or increased FA synthesis through DNL in this primitive population. HPC1/2s, on the other hand, were enriched for carnitine containing lipid metabolites (**Figure 1A**). This suggests increased utilization of FAs for mitochondrial metabolism in progenitor populations downstream of HSC/MPPs.

**Figure 1.**
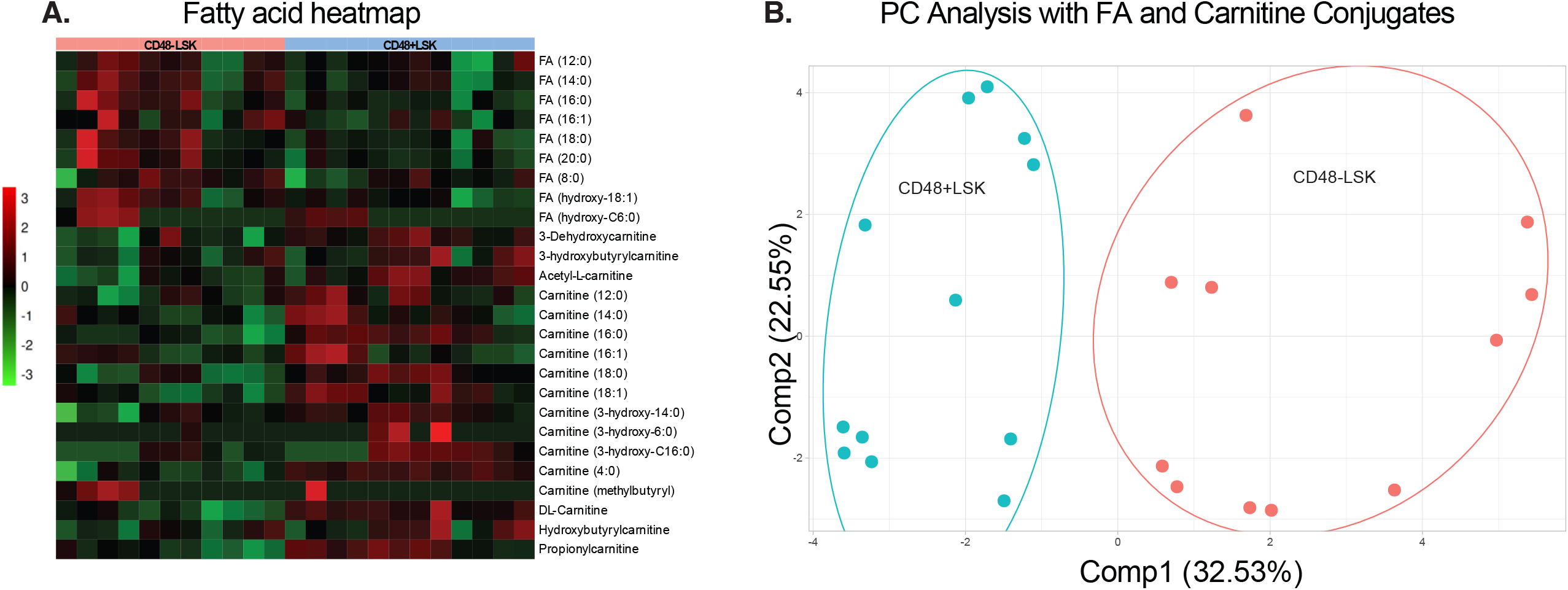
Hematopoietic stem and primitive progenitor cells maintain a fatty acid pool.

Principal component analysis (PCA) based on FAs and carnitine conjugates demonstrated distinct clustering of HSC/MPPs and HPC1/2s (**Figure 1B**). PC1 Collectively, these data indicate that while FAs in downstream progenitor cells are more likely to have features characteristic of mitochondrial FA metabolism, primitive HSC/MPPs maintain a FA pool, the source of which is unknown.

### Steady state hematopoiesis is dependent on DNL through ACC1

To investigate whether or not hematopoietic stem cells are dependent on DNL, we utilized a previously described floxed *Acaca* allele (encoding ACC1) crossed to mice harboring *Mx1Cre*.^11,12^ This strategy facilitated deletion of *Acaca* in homozygous *Acaca*^*fl/fl*^ *Mx1Cre+* (henceforth KO) mice following 5 every other day polyinosinic:polycytidylic acid (poly (I:C)) injections (**Figure S1A-B**). Within 6 weeks of deletion, KO mice demonstrated slight decreases in white blood cell count and hemoglobin, but more profound thrombocytopenia by complete blood counts (**Figure 2A**). KO mice also demonstrated splenomegaly (**Figure 2B, 2D, and 2E**) and reduced bone marrow cellularity (**Figure 2C**). Flow cytometric analysis of the peripheral blood and spleen demonstrated a relative myeloid expansion and B cell lymphopenia (**Figure 2F and 2G**). Comparable analysis of the bone marrow did not reveal significant differences in the mature myeloid or lymphocyte populations (**Figure 2H**). Immunophenotypic analysis of the common myeloid progenitor, megakaryocyte-erythrocyte progenitor, and granulocyte-monocyte progenitor compartments did not demonstrate significant differences (**Figure S1C**). These data demonstrate that intact ACC1 is required to maintain hematopoietic homeostasis.

**Figure 2.**
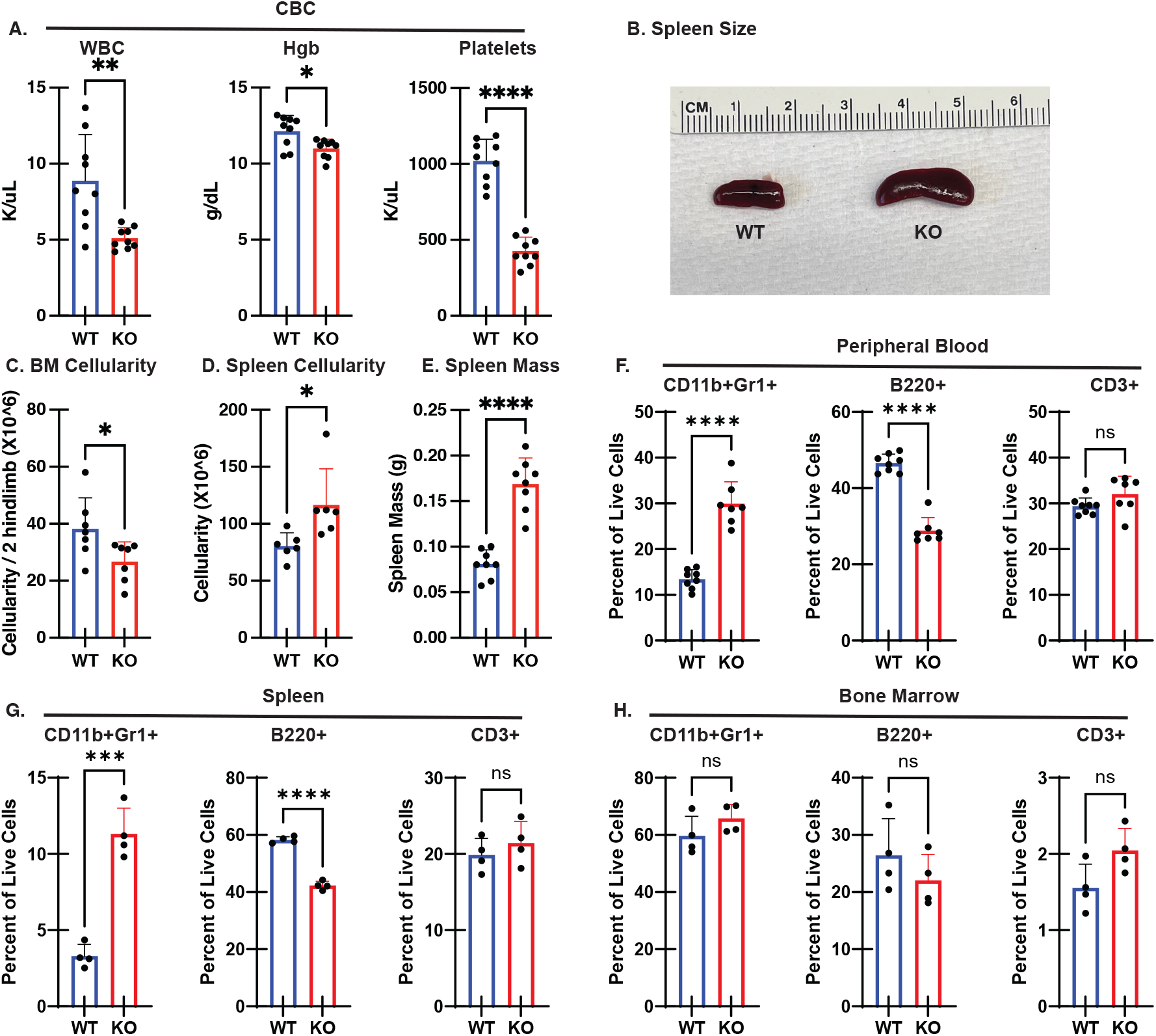
ACC1 loss impacts steady state hematopoiesis 6-8 week after gene inactivation. (A) Complete blood count evaluation of white blood cell count (WBC), hemoglobin (Hgb), and platelets in wildtype (WT) and ACC1 knockout (KO) mice evaluated 6-8 weeks follows poly (I:C) injection. (B) Representative photo of WT and KO spleens demonstrating splenomegaly in KO mice. (C) Bone marrow (BM) cellularity comparing WT and KO. (D and E) Spleen cellularity (D) and mass (E) comparing WT and KO. (F-H) Flow cytometric evaluation of myeloid (CD11b+Gr1+), B cells (B220+), and T cells (CD3+) in the peripheral blood (F), spleen (G), and bone marrow (H) of WT and KO mice. In the above figure data are derived from n = 4-9 individual mice per group with values from individual mice shown. Graphs display mean + S.D. Student’s t test was used for comparison of means. P values *<0.05, **<0.01, ***<0.001, ****<0.0001.

To immunophenotypically characterize the stem cell compartment in WT and KO mice 6 weeks after *Acaca* deletion, the SLAM stain was applied to the bone marrow.^13^ This revealed a marked expansion of the KO Lineage-cKit+Sca1+ (LSK) population (**Figure 3A,B,F**). This compartment includes the long-term hematopoietic stem cell (CD48-CD150+LSK; LT-HSC), multipotent progenitor (MPP; CD48-CD150-LSK), and hematopoietic progenitor 1/2 (HPC1/2; CD48+LSK) populations (**Figure 3A, C-E, G-I**). Analysis demonstrated that the most prevalent population expansion was detected in the KO LT-HSCs and KO HPC1/2s. Spleens from WT and KO mice were also subjected to the SLAM stain. This revealed an atypical expansion of LSK, including LT-HSCs, in KO mice, indicative of extramedullary hematopoiesis (**Figure 3J-R**). The data indicate that the loss of ACC1 results in the expansion of immunophenotypically-defined primitive hematopoietic populations and aberrant splenic mobilization.

**Figure 3.**
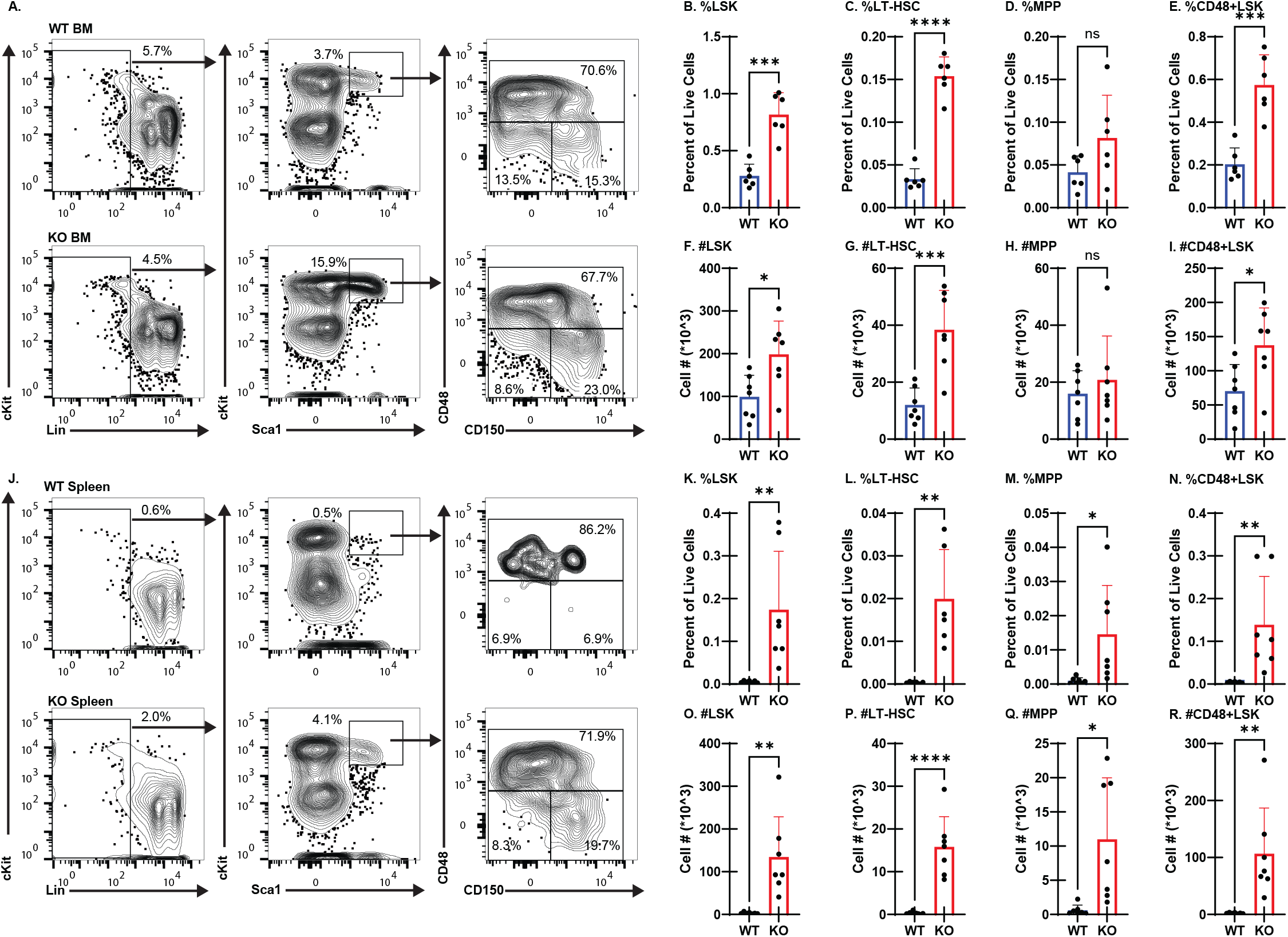
ACC1 loss results in expansion of hematopoietic stem and primitive progenitor cells in the bone marrow and spleen. (A) Representative flow cytometric gating strategy of WT and KO bone marrow 6-8 weeks after poly (I:C) injection. (B-E) Frequency of LSK (Lineage^-^Sca-1^+^cKit^hi^), LT-HSC (CD48^-^CD150^+^LSK), MPP (CD48^-^CD150^-^LSK), and HPC1/2 (CD48^+^LSK) in the bone marrow. (F-I) Absolute numbers of LSK, LT-HSC, MPP, and HPC1/2 populations in the bone marrow. (J) Representative flow cytometric gating strategy of WT and KO spleens 6-8 weeks after poly (I:C) injection. (K-N) Frequency of LSK, LT-HSC, MPP, and HPC1/2 in the spleen. (O-R) Absolute numbers of LSK, LT-HSC, MPP, and HPC1/2 in the spleen. In the above figure data are derived from n = 6-7 individual mice per group with values from individual mice shown. Graphs display mean + S.D. Student’s t test was used for comparison of means. P values *<0.05, **<0.01, ***<0.001, ****<0.0001.

To determine if the impact of *Acaca* loss on steady state hematopoiesis was specific to the *Mx1Cre* driver and/or the inflammatory impact of poly (I:C) injection, we crossed the *Acaca*^*fl*^ mice to mice harboring *Vav1Cre*.^14^ *Vav1Cre* is activated at embryonic day 11.5 in hematopoietic cells. This strategy did not yield any live births. We therefore analyzed embryos at e15.5-e16.5. KO embryos demonstrated pallor relative to WT (**Figure S2A**). No significant differences in fetal liver cellularity, myeloid cells, B cells, or T cells were observed (**Figure S2B-D**). Similar to the *Mx1Cre* model, the KO LSK progenitor compartment, including the LT-HSCs, was markedly expanded when *Acaca* was deleted via *Vav1Cre* (**Figure S2E-H**). Mice that lost a single allele of *Acaca* did not have any detectable phenotype. This indicates that hematopoietic progenitor expansion is neither specific to *Mx1Cre* nor the result of poly (I:C) induced inflammation.

### ACC1 loss impairs HSC function in bone marrow transplantation

To determine if ACC1 loss impacts HSC function despite immunophenotypic expansion, bone marrow transplantation assays were established. 500,000 mononuclear cells from the bone marrow of WT or KO mice were mixed in a 1:1 ratio with B6.SJL competitor cells **(Figure 4A**). Following transplantation into lethally irradiated recipients, engraftment was measured via flow cytometry. After documentation of engraftment, 5 every other day injections of poly (I:C) were administered to activate Mx1Cre and thus inactivate ACC1. Myeloid (CD11b+Gr1+), B cell (B220+), and T cell (CD3+) engraftment was then measured every 4 weeks in the peripheral blood via flow cytometry (**Figure 4B-C**). These data demonstrate a profound defect in trilineage hematopoietic output from the KO tester graft. Since baseline engraftment was documented and ACC1 deletion occurred following engraftment, this defect cannot be explained by impaired KO HSC homing.

**Figure 4.**
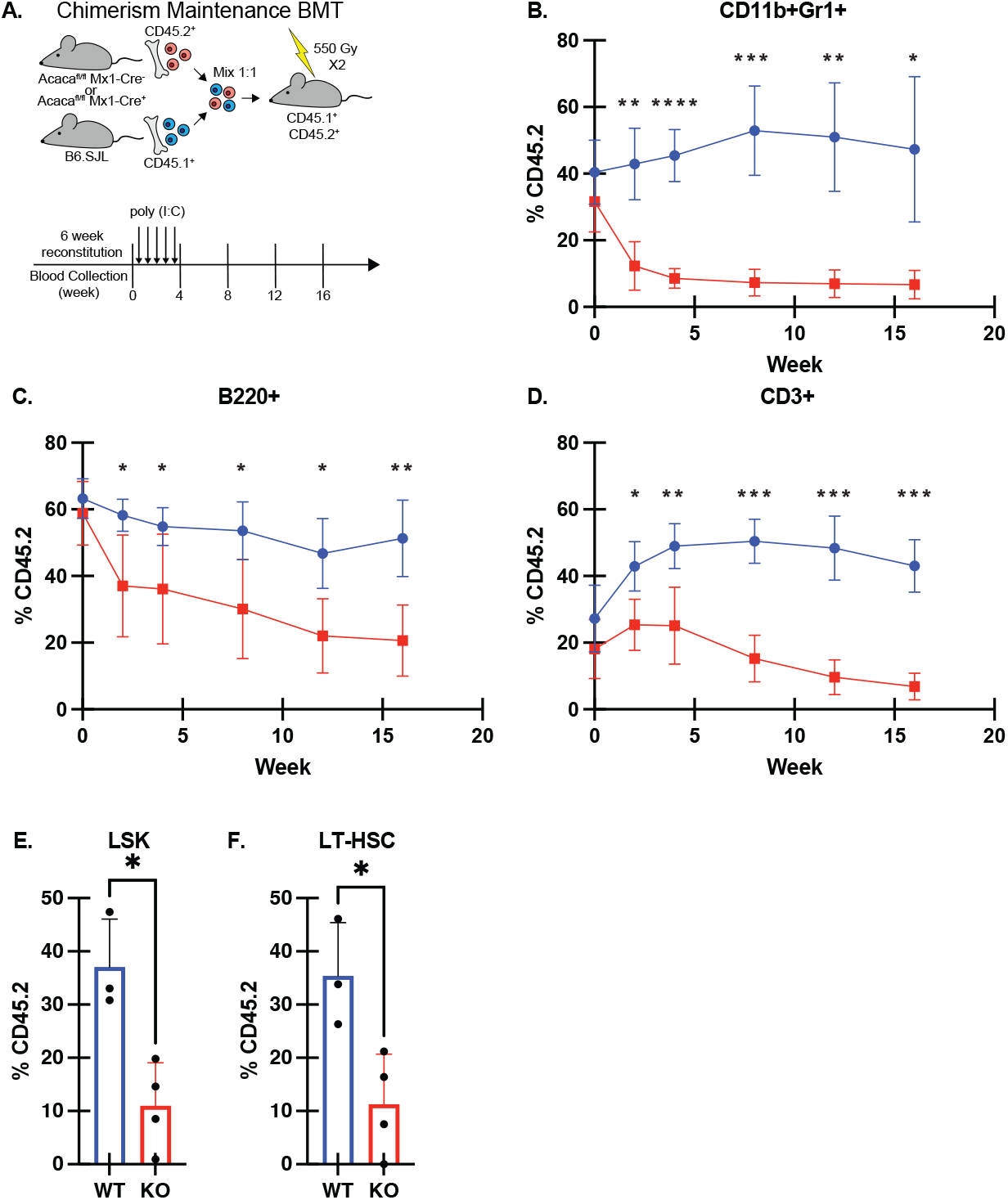
ACC1 loss results in functional defects in hematopoietic stem cells in bone marrow transplantation assays. (A) Experimental design demonstrating competitive boné marrow transplantation approach in which bone marrow chimeras were generated with CD45.2^+^WT or KO competitor bone marrow mixed 1:1 with CD45.1^+^ tester bone marrow prior to deletion of ACC1. Engraftment was documented by flow cytometry prior to poly (I:C) injection into both WT and KO graft recipients. Engraftment was then measured in the peripheral blood at indicated timepoints and transplant was terminated after 16 weeks. (B) Myeloid (CD11b^+^Gr1^+^) output in peripheral blood at indicated timepoints measured by flow cytometry. (C) B cell (B220^+^) output in peripheral blood at indicated timepoints measured by flow cytometry. (D) T cell (CD3^+^) output in peripheral blood at indicated timepoints measured by flow cytometry. (E-F) Donor LSK and LT-HSC frequency in bone marrow after 16 weeks as measured by flow cytometry. In the above figure data are derived from n = 3 individual mice per group with values from individual mice shown. Graphs display mean + S.D. Student’s t test was used for comparison of means. P values *<0.05, **<0.01, ***<0.001, ****<0.0001.

After 16 weeks of monitoring, recipient mice were sacrificed and the LSK and LT-HSC compartments were analyzed by flow cytometry. Analysis of both compartments demonstrated hematopoietic stem and progenitor depletion in KO graft recipients, indicating that defective function was tractable to LT-HSCs (**Figure 4E-F**).

Collectively, these data demonstrate that despite immunophenotypic expansion in steady state conditions, KO HSCs show reduced capacity of 1) maintaining trilineage hematopoietic reconstitution and 2) maintaining self-renewal in bone marrow transplantation assays. Thus, ACC1 is required for the maintenance of a functional LT-HSC pool.

### ACC1 loss drives a loss of quiescence in LT-HSCs

Preserved HSC function requires that LT-HSCs be maintained in a largely quiescent state. An inability to maintain quiescence has been shown to lead to progressive LT-HSC functional decline in multiple models.^15-20^ To investigate if KO LT-HSCs maintained quiescence, WT or KO mice were injected with 5-ethynyl-2’-deoxyuridine (EDU) and then placed on EDU containing water for 72hrs. Flow cytometric analysis demonstrated that the KO LT-HSC pool incorporated more EDU than WT LT-HSCs (**Figure 5B**). This was not true in the overall LSK population (**Figure 5A**). This suggests that KO LT-HSCs are more proliferative than WT.

**Figure 5.**
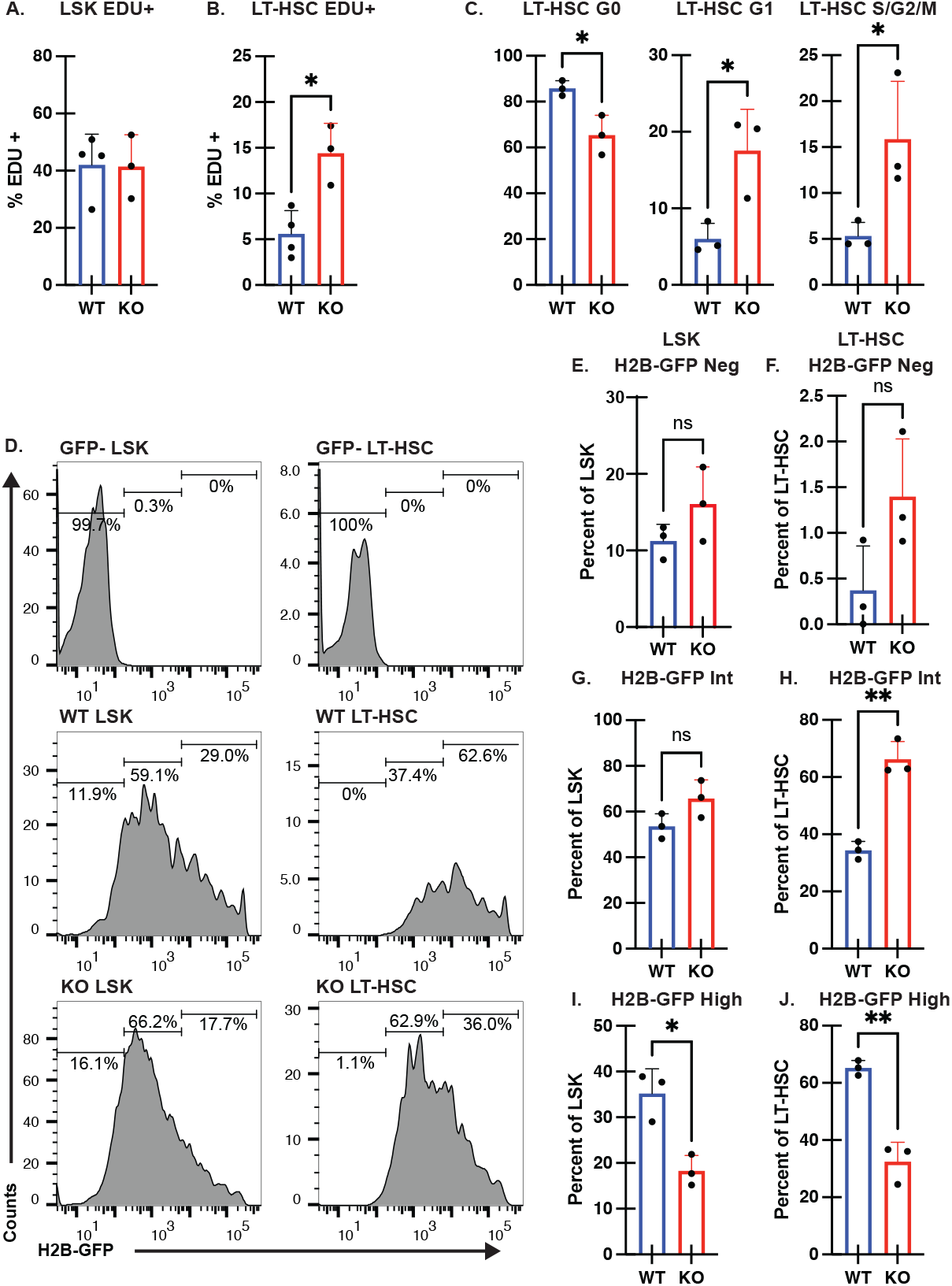
ACC1 is required for the maintenance of HSC quiescence. (A-B) Comparison of EDU uptake WT or KO LSK (A) and LT-HSC (B) compartments following 1mg EDU IP injection and 72hr labeling period with EDU water(???). (C) Cell cycle analysis by Ki67 staining demonstrating reduced LT-HSCs in G0 and increase in G1/S/G2/M in KO as compared to WT. (D) Schematic representation of H2B-GFP pulse-chase experiment. (E) Representative flow cytometry histograms of H2B-GFP distribution in WT and KO LSK and LT-HSC following 12-week chase period. (F-K) Comparison of means of percentage of cells in H2B-GFP negative (Neg), intermediate (int), and high fractions in LSK (F,H, J) and LT-HSC (G, I, K) populations. In the above figure data are derived from n = 3 individual mice per group with values from individual mice shown. Graphs display mean + S.D. Student’s t test was used for comparison of means. P values *<0.05, **<0.01.

To better assess the quiescent, G0 state, Ki67 staining was also used to analyze cell cycle status. Using this strategy, fewer KO LT-HSCs were found to be in G0 and were proportionally identified in later phases of the cell cycle compared to WT (**Figure 5C**). This demonstrated that KO LT-HSCs failed to maintain an essential quiescent state.

EDU and Ki67 staining provide snapshots into cell cycle activity at defined timepoints. To determine if increased cell cycle activity was sustained over time in the KO LT-HSC population, we crossed KO mice to a previously described tetracycline activated *H2bGFP* allele along with *M2rTTA* as previously described.^16,21^ We then activated H2B-GFP expression with doxycycline during a 6-week pulse and analyzed the LSK and LT-HSC compartment after a subsequent 12-week chase period following removal of doxycycline. This revealed that KO LT-HSCs diluted H2B-GFP much faster than WT LT-HSCs (**Figure 5D,F, H,J**). The KO LSKs also diluted H2B-GFP faster than WT LSK, but the effect was not as profound as that seen in the LT-HSCs (**Figure 5D, E, G, I**). The loss of ACC1, therefore, results in an inability for LT-HSCs to maintain or revert to a quiescent phenotype.

### ACC1 loss results in alteration in transport of extracellular FAs and increased oxidative stress

Since ACC1 is the first and rate limiting step of DNL, we hypothesized that the loss of DNL capacity would require that hematopoietic stem and progenitor cells become dependent on external FFAs for cellular needs. To measure this, we used C12 BODIPY, a fluorescent construct that resembles a 16-carbon fatty acid and can be taken up into cells that have the capacity to transport extracellular FAs to internal compartments. In so doing, we found that KO LT-HSCs and LSK progenitors internalized more C12 BODIPY than WT counterparts, suggesting increased fat uptake upon loss of ACC1 (**Figure 6A,B**).

**Figure 6.**
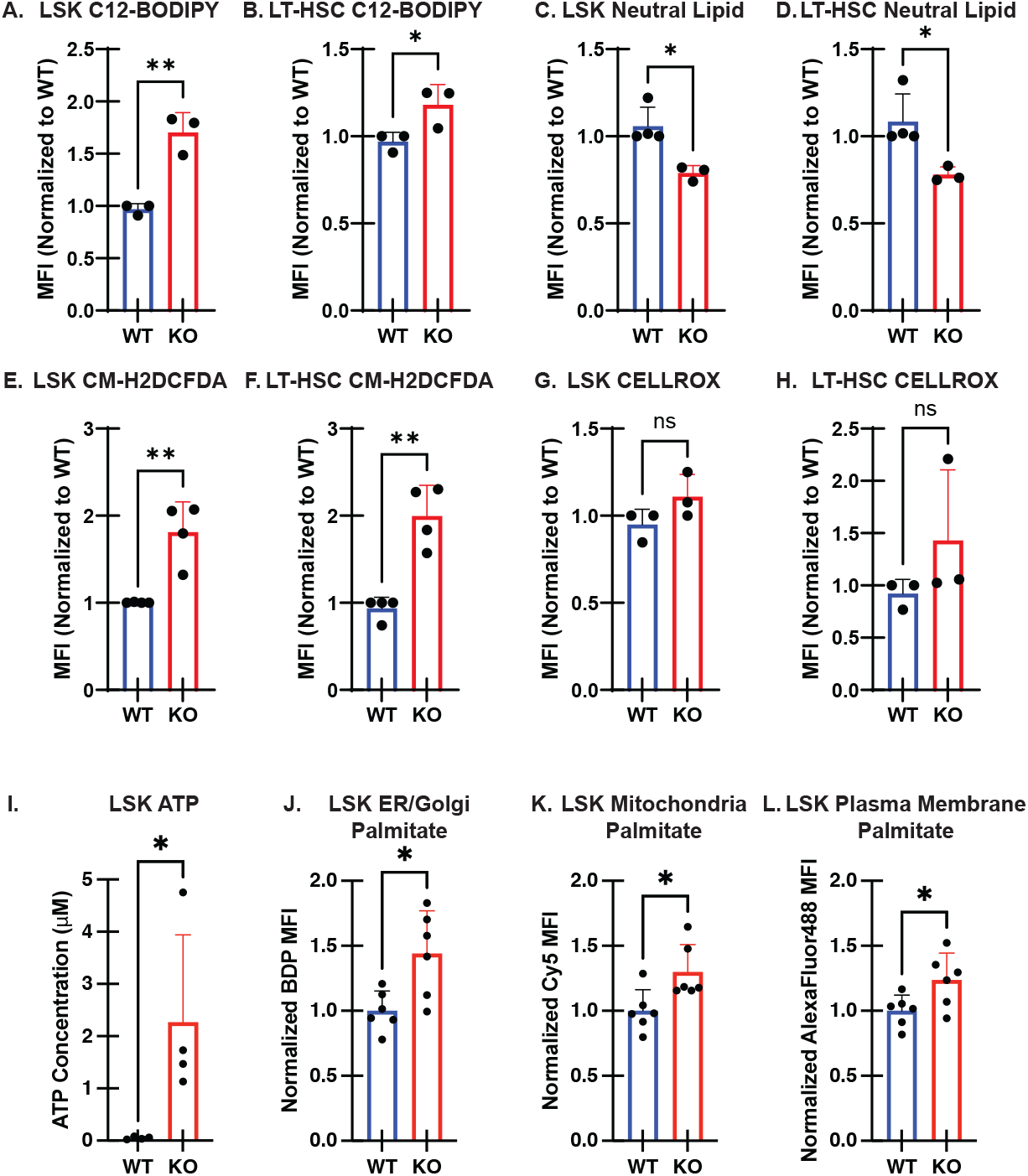
Loss of ACC1 results in increased uptake of extracellular fatty acids and increased mitochondrial activation. (A) Flow cytometric evaluation of C12-BODIPY uptake over 30 minutes in LSK in WT and KO mice. (B) Flow cytometric evaluation of C12-BODIPY uptake over 30 minutes in LT-HSC in WT and KO mice. (C) Flow cytometric evaluation of neutral lipid content using BODIPY 493/503 staining in LSK in WT and KO mice. (D) Flow cytometric evaluation of neutral lipid content using BODIPY 493/503 staining in LT-HSC in WT and KO mice. (E) Flow cytometric identification of Palmitate-Alkyne incorporation into ER/Golgi membrane using modified O-CLICKFC in LSK of WT and KO mice. (F) Flow cytometric identification of Palmitate-Alkyne incorporation into the mitochondria using modified O-CLICKFC in LSK of WT and KO mice. (G) Flow cytometric identification of Palmitate-Alkyne incorporation into the plasma membrane using modified O-CLICKFC in LSK of WT and KO mice. (H) Flow cytometric evaluation of ROS in LSK using CM-H2DCFDA in WT and KO mice. (I) Flow cytometric evaluation of ROS in LT-HSC using CM-H2DCFDA in WT and KO mice. (J) ATP concentration in LSK of WT and KO mice using Roche ATP bioluminescence kit. In the above figure data are derived from n = 3-5 individual mice per group with values from individual mice shown. Graphs display mean + S.D. Student’s t test was used for comparison of means. P values *<0.05, **<0.01.

After uptake, FAs can be utilized for bioenergetic needs, as biomass, and/or stored in lipid droplets. To determine if extracellular fat was accumulating in lipid droplets in KO mice, BODIPY neutral lipid (NL) staining was utilized. NL staining demonstrated that KO LSK and LT-HSCs had less NL content than WT, suggesting that the increase in external FA uptake did not lead to NL accumulation in lipid droplets in KO hematopoietic stem and progenitor cells (**Figure 6C,D**).

β-oxidation is the process in which fat is catabolized in the mitochondria to be used as fuel and is associated with an increase in oxidative phosphorylation in HSCs. To achieve this, FAs must be transported into the mitochondria. This process is highly regulated plays essential roles in HSC function.^4,7^ To better characterize compartmental localization of FA in cells following uptake, we adapted a previously described strategy termed organelle-selective click labeling coupled with flow cytometry (O-ClickFC).^22^ In this strategy, a “clickable” substrate is fed to cells *ex vivo* and then through utilization of fluors with selective permeability, localization to the ER/golgi, mitochondria, and plasma membrane can be identified. We exposed WT or KO bone marrow cells to alkyne labeled palmitate and then identified localization. We found that this substrate was distributed throughout the ER/golgi, mitochondria, and plasma membrane of KO LSK progenitors to an increased degree as compared to WT (**Figure 6J-L**). This indicates that extracellular FAs are being internalized and processed with distribution throughout KO progenitors, including trafficking to the mitochondria.

We hypothesized that if extracellular FAs were indeed being directed to the mitochondria to be used for energy, HSPCs would demonstrate hallmarks of increased oxidative metabolism. Indeed, KO LT-HSCs and LSKs demonstrated an increase in the production of reactive oxygen species as demonstrated by CM-H2DCFDA staining, but not Cellrox (**Figure 6E-H**). Additionally, there was a concomitant increase in ATP produced in KO LSK progenitor cells (**Figure 6I**). Together, these data suggest that extracellular FA utilization and increased mitochondrial β-oxidation may be increased in KO HSPC population, including LT-HSCs.

### CD36 loss improves ACC1-deficient hematopoiesis

CD36 has previously been shown to be required in HSPCs for extracellular uptake of FAs to fuel mitochondrial β oxidation under infectious stress.^1^ To date, this has been the only FA transporter suggested to be active in primitive hematopoietic progenitors. Increased extracellular fat utilization and concomitant mitochondrial activation has been demonstrated to be a mechanism of HSC dysfunction under high fat feeding conditions.^7^ We therefore reasoned that if increased FA uptake was responsible for phenotype observed in KO animals, deletion of CD36 may abrogate this. To determine if CD36 inhibition improved ACC1-deficient hematopoiesis, we bred mice homozygous for a previously described floxed CD36 allele^23^ to our ACC1 floxed animals to generate mice with the *Cd36*^*fl/fl*^ *Acaca*^*fl/fl*^ *Mx1Cre*^*+*^ genotype, along with requisite control animals. We then characterized steady state hematopoiesis 4-6 weeks after completion of 5 every other day poly (I:C) injections.

We found that animals with the combined loss of ACC1 and CD36 had reduced splenomegaly compared to those with the loss of ACC1 alone (**Figure 7A**). The loss of CD36 alone did not impact spleen size. Following loss of ACC and CD36, peripheral blood analysis demonstrated a significant improvement in thrombocytopenia, as compared to the ACC1 single knockout, and did not cause significant changes in WBC or Hgb (**Figure 7B-D**). Additionally, these combined knockout animals had reduced myeloid skewing in the spleen (**Figure 7E**). CD36 knockout alone did not have any impact on these steady state hematologic parameters. Together, these data indicate that the loss of CD36 improves elements of the ACC1 KO phenotype downstream of hematopoietic stem and progenitor cells.

**Figure 7.**
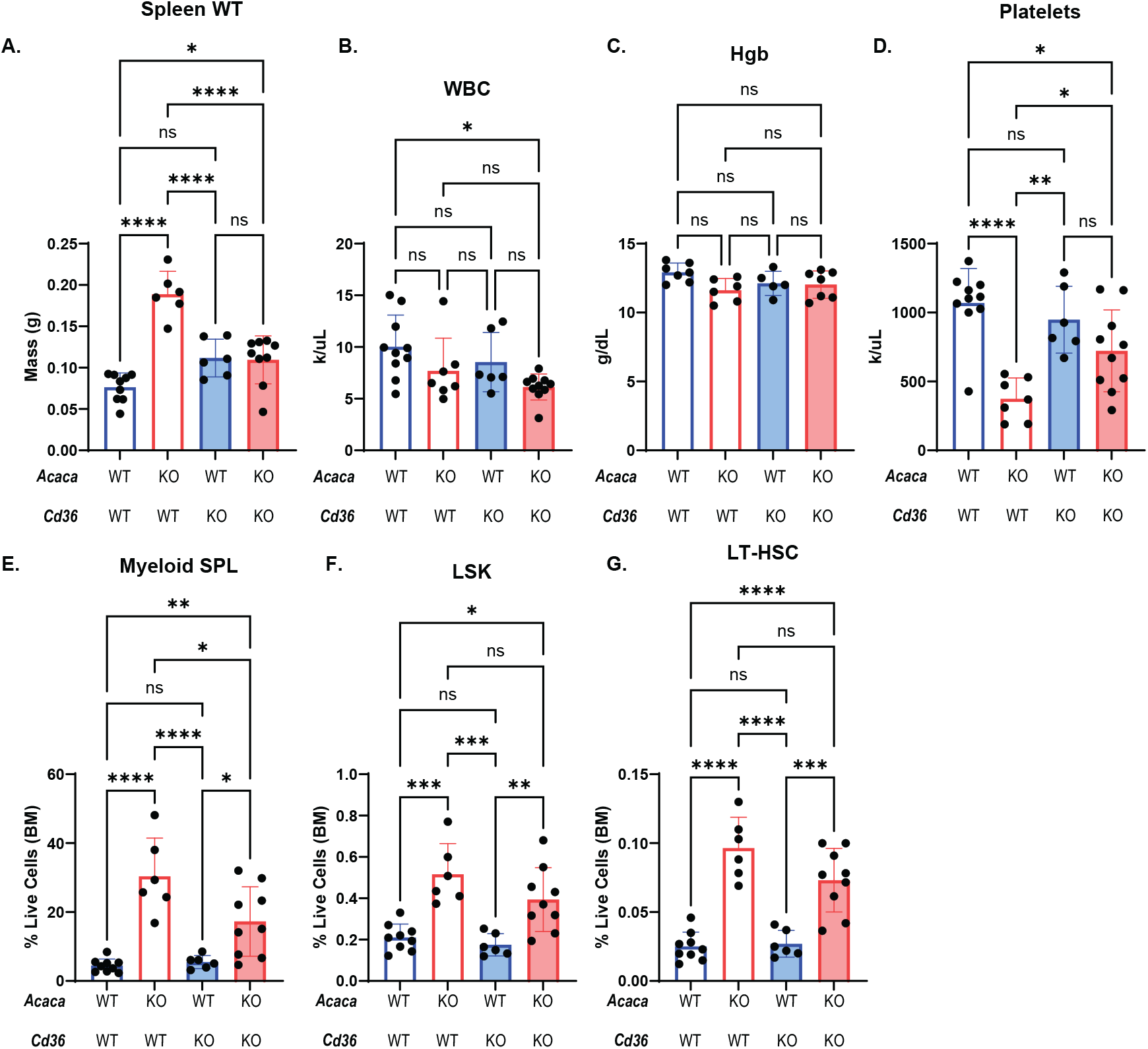
CD36 deficiency does not correct the ACC1 LT-HSC phenotype but improves downstream hematopoiesis. (A) Comparison of spleen weights between WT, ACC1 KO, CD36 KO, and ACC1 CD36 DKO mice. (B-D) CBC measurement of WBC, Hgb, and platelets in WT, ACC1 KO, CD36 KO, and ACC1 CD36 DKO mice. (E) Comparison of myeloid (CD11b+Gr1+) population in the spleen of WT, ACC1 KO, CD36 KO, and ACC1 CD36 DKO mice by flow cytometry. (F) Comparison of frequency of LSK (Lineage^-^Sca-1^+^cKit^hi^) population in the bone marrow of WT, ACC1 KO, CD36 KO, and ACC1 CD36 DKO mice by flow cytometry. (G) Comparison of frequency of LT-HSC (CD48^-^CD150^+^LSK) population in the bone marrow of WT, ACC1 KO, CD36 KO, and ACC1 CD36 DKO mice by flow cytometry. In the above figure data are derived from n = 6-10 individual mice per group with values from individual mice shown. Graphs display mean + S.D. Anova was used for comparison of means. P values *<0.05, **<0.01, ***<0.001, ****<0.0001.

We next evaluated the hematopoietic stem and progenitor compartment of combined knockout animals. This demonstrated that unlike the peripheral blood and mature myeloid compartments, the combination of CD36 loss with ACC1 loss did not impact the LSK and LT-HSC expansion phenotype identified in the ACC1 KO animals (**Figure 7F-G**). CD36 loss alone did not have an impact on the LSK and HSC compartments.

Collectively, these data thus indicate that FA uptake through CD36 is not directly responsible for the LT-HSC expansion observed in ACC1 KO animals but may play a role in progenitors downstream of LT-HSCs.

### Cpt1a deletion partially rescues ACC1-deficient hematopoietic reconstitution

CPT1A serves as the so-called gate keeper to mitochondrial β–oxidation and generates acyl carnitines from fatty acyl-CoAs and carnitine to facilitate transport into the mitochondria. CPT1A loss has been shown to improve HSC function when mitochondrial β-oxidation is excessively engaged through high fat feeding.^7^ We therefore reasoned that increased FA uptake, ATP generation, and ROS accumulation in ACC1 KO hematopoietic stem and progenitor cells could be the result enhanced β–oxidation in our model and thus LT-HSC defects could be rescued through preventing β–oxidation activation.

To achieve this, *Acaca*^*f/f*^*Mx1-Cre*^*+*^ mice were crossed with previously described mice harboring the conditional *Cpt1a*^*f*^ to generate mice with the *Acc1*^*f/f*^*Cpt1a*^*f/f*^*Mx1Cre*^*+*^ genotype. WT, *Acaca*^*f/f*^*Mx1-Cre*^*+*^ (KO), and *Acc1*^*f/f*^*Cpt1a*^*f/f*^*Mx1Cre*^*+*^ (DKO) transplant donor animals were injected with 3 doses of Poly (I:C). 500,000 whole bone marrow cells from each donor were then mixed with 500,000 congenic CD45.1+ competitor cells and injected into lethally irradiated recipients (**Figure 8A**). Throughout 20 weeks of peripheral blood monitoring, no differences were identified in output into the CD11b+Gr1+ (myeloid), B220+ (B cell), and CD3+ (T cell) populations (**Figure 8B-D**).

**Figure 8.**
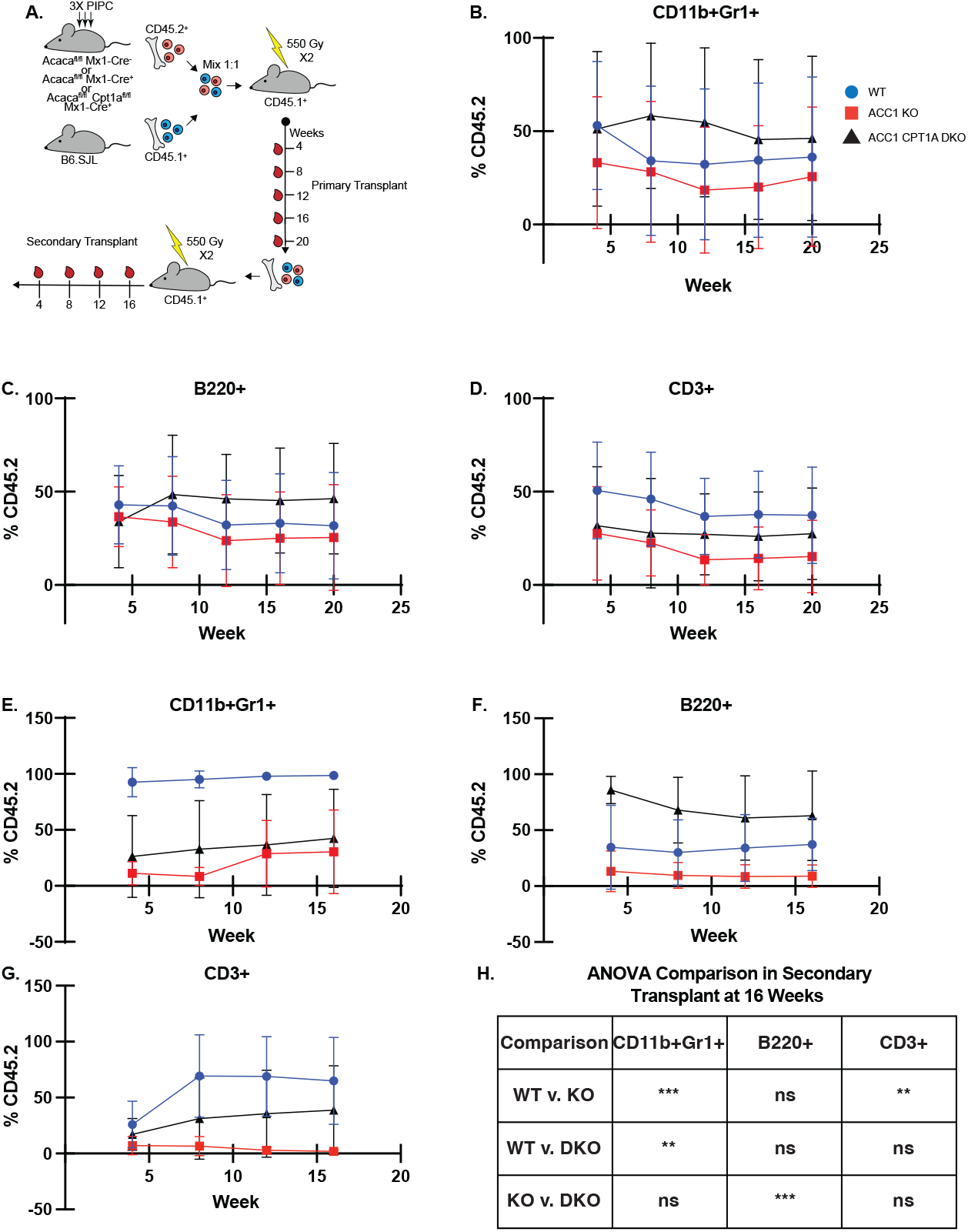
CPT1A loss partially rescues hematopoietic reconstitution in ACC1 KO transplantation models. (A) Experimental design demonstrating competitive bone marrow transplantation approach in which bone marrow chimeras were generated with CD45.2^+^WT, ACC1 KO, or ACC1 CPT1A DKO tester bone marrow mixed 1:1 with CD45.1^+^ competitor bone marrow. Donor mice were treated with 3 doses of poly (I:C) prior to transplant. Engraftment was then measured in the peripheral blood at indicated timepoints and transplant was terminated after 20 weeks. Bone marrow from a subset of recipients was then harvested an retransplanted to establish secondary bone marrow transplants. Engraftment was again measured at the indicated time points. (B) Myeloid (CD11b^+^Gr1^+^) output in peripheral blood at indicated timepoints measured by flow cytometry in primary transplants. (C) B cell (B220^+^) output in peripheral blood at indicated timepoints measured by flow cytometry in primary transplants. (D) T cell (CD3^+^) output in peripheral blood at indicated timepoints measured by flow cyt
ometry in primary transplants. (E) Myeloid (CD11b^+^Gr1^+^) output in peripheral blood at indicated timepoints measured by flow cytometry in secondary transplants. (F) B cell (B220^+^) output in peripheral blood at indicated timepoints measured by flow cytometry in secondary transplants. (G) T cell (CD3^+^) output in peripheral blood at indicated timepoints measured by flow cytometry in secondary transplants. (H) Statistical comparison of donor contribution in specified lineages at 16 weeks post transplant by Anova. In the above figure data are derived from n >/= 7 individual transplant recipient mice per group. Graphs display mean +/-S.D. Anova was used for comparison of means. Significant P values reported in (H). No statistical significance was identified elsewhere. **<0.01, ***<0.001.

After the 20-week primary transplant experiment, total bone marrow from a subset of recipient animals was transplanted into new lethally irradiated recipient mice to establish secondary transplants (**Figure 8A**). WT donor bone marrow robustly maintained myeloid output while KO and DKO bone marrow were less effective in this regard (**Figure 8E**). WT engraftment in the myeloid lineage was significantly higher than both KO and DKO, and no difference was observed between KO and DKO at 16 weeks (**Figure 8H**). In the B cell linage, there was apparent, but not statistically significant differences detected between WT and KO output and WT and DKO output. Intriguingly, DKO B cell output was increased compared to both WT and KO (**Figures 8F, H**). In the T cell lineage, T cell output from the WT and DKO donor bone marrow was increased compared to KO, with only the difference between WT and KO achieving statistical significance (**Figure 8G, H**). Taken together, these data suggest that increased β–oxidation may impact hematopoietic reconstitution following ACC1 inactivation, but it does not fully rescue the trilineage hematopoietic defects identified in ACC1-deficient HSCs.

## Discussion

While oxidative fatty acid metabolism has been demonstrated as both essential to hematopoietic stem cell function and detrimental under times of stress, fatty acid anabolic metabolism has not been characterized in hematopoietic stem cells.^4,7^ This report represents the first study demonstrating that intact *de novo* lipogenesis is essential to HSC function.

We found that while downstream hematopoietic progenitor cells have a relative increase in acyl carnitines, signifying increased β oxidation potential, primitive hematopoietic stem cells maintain a pool of fatty acids. The source of this pool of fatty acids was unknown. Extracellular fatty acids appear to be used in times of stress^1^, but endogenous synthesis through *de novo* lipogenesis had not been demonstrated as active and therefore had not been investigated as a contributor to this pool. In deleting ACC1, the first and rate-limiting step of *de novo* lipogenesis, we demonstrate that intact DNL through ACC1 is necessary for hematopoietic homeostasis. The development of thrombocytopenia, splenomegaly, extramedullary hematopoiesis, myeloid skewing, and HSC expansion phenotypically resembled common features of myeloid neoplasms.^24-27^ Neoplastic HSCs, however, often have increased self-renewal capacity compared to normal HSCs. The ACC1-deficient HSC population, though expanded, is functionally impaired. ACC1-deficient HSCs fail to support trilineage hematopoietic output and maintain engraftment in transplantation experiments. These data collectively indicate that ACC1 is required to restrain hematopoiesis and maintain balanced hematopoietic output in steady state. The data further suggest that long-term self-renewal requires DNL through ACC1.

Ultimately, DNL through ACC1 is required to maintain HSC quiescence. ACC1-deficient HSCs do not maintain quiescence and instead engage in both short term and sustained cell cycle activity, as demonstrated by Ki67/EdU and H2B-GFP pulse-chase experiments, respectively. An inability to maintain quiescence has been shown to be detrimental to HSC function.^16-20^ While this finding could be a compensatory effect as HSCs attempt to generate progenitors to account for thrombocytopenia, several key findings argue against this. First, megakaryocyte-erythrocyte Progenitors (MEPs) are not expanded in KO hematopoietic tissues. This would be expected if there was bone marrow compensation for thrombocytopenia, unless a differentiation block existed between the HSPC and MEP developmental stages. Second, our experiments with combined ACC1 and CD36 deficient hematopoiesis demonstrate hematopoietic stem and progenitor expansion despite the improvement in platelet counts. Instead, we hypothesize that a loss of ACC1 and a dependence on extracellular FAs drives this loss of quiescence. Indeed, several studies assessing HSC behavior subjected to high fat diet feeding demonstrate damage to the HSC pool and a loss of quiescence.^28,29 30^ The mechanisms through which steady state HSCs import extracellular FAs and the utilization of extracellular FAs remain under further investigation.

A proposed mechanism through which enhanced extracellular fatty acids damage hematopoietic stem cells is enhanced mitochondrial β-oxidation. ACC1-deficient hematopoietic stem and progenitor cells have increased ATP, ROS and fatty acid uptake. All of this led to the hypothesis that this enhanced β-oxidation accounted for HSC functional deficits in our ACC1 model. CD36 has been described as modulating extracellular fatty acid uptake and β-oxidation under infectious stress.^1^ We reasoned that inactivating CD36 could therefore abrogate HSC stress related to increased fatty acid uptake/mitochondrial β-oxidation in ACC1-deficient hematopoiesis. We found that while splenomegaly was diminished and thrombocytopenia was improved, immunophenotypic expansion of HSCs persisted. This suggests that CD36-deficiency improved features of hematopoiesis downstream of HSCs, but did not contribute to HSC expansion.

We then sought to inactivate β-oxidation completely through conditional inactivation of CPT1A in ACC1-deficient hematopoiesis. This provided a more direct means of determining if oxidative fatty acid metabolism was detrimental to our ACC1 KO hematopoietic stem cells. Using this approach, we found that CPT1A deficiency partially rescued hematopoietic reconstitution in ACC1-deficient hematopoietic stem cells. This demonstrates that the impact of ACC1 deficiency on hematopoietic stem cell function cannot be fully explained by increased mitochondrial β-oxidation.

Our report demonstrates an essential and multifaceted role for *de novo* lipogenesis through ACC1 in HSC function. This role extends beyond hyperactivate β-oxidation in ACC1-deficient HSCs but implicates DNL through ACC1 as a critical regulatory of HSC quiescence. Further lipidomic studies addressing the impact of ACC1 loss on intercellular lipid species are necessary next steps in understanding how specific lipids impact hematopoietic stem cell homeostasis. This work will ultimately provide further insight into how HSC obtain and utilize fatty acids and the broad impact that this has on HSC function.

## EXPERIMENTAL MODEL DETAILS

*Acaca*^*fl*^, *Cpt1a*^*fl*^, and *Mx1-Cre* mice were obtained from Jackson Laboratories.^11,12,31^ Mice were all maintained on the C57BL/Ka-CD45.2:Thy1.1 background following a minimum backcross of 10 generations. Cre expression in *Mx1-Cre* mice was induced via injection of polyinosinic:polycytidylic acid (poly I:C) dissolved in PBS on an every other day intraperitoneal injection schedule for a total of 3-5 injections once mice were 6-8 weeks of age. Transplantation experiments utilized C57BL/Ka-CD45.1:Thy1.2 and C57BL/Ka-CD45.2:Thy1.1X C57BL/Ka-CD45.1:Thy1.2 (F1) mice as competitor bone marrow source and recipients as annotated. Fetal liver studies were performed in 15.5-16.5 days post coitum pregnant females. No differences were identified between mouse genders and therefore both genders were pooled for data analysis. Equal numbers of male and female mice were used where able. For experiments where doxycycline was used to activate H2B-GFP expression through the TET operator, doxycycline was added to water at a concentration of 0.2%(m/V) along with 1% sucrose. Animals were maintained on water for 6 weeks prior to withdrawal.

## METHOD DETAILS

### Hematopoietic cell isolation and flow cytometry

Bone marrow cells were isolated either through flushing of the bone marrow from femurs and tibias or crushing of the spine, hips, femurs and tibias (when increased cell number was desired). Flushing was performed using a syringe affixed with a 25 gauge needle and FACS buffer (Hanks buffer without calcium or magnesium and with 2% heat inactivated calf serum). Cell suspension was then mixed using the same syringe before being passaged through a 70μM filter mesh to generate a single cell suspension. Crushing was performed using a mortar and pestle with FACS buffer maintained on ice. Dissociation was achieved by pipetting crushed mixture via pipet prior to passaging through a 70μM filter mesh to generate a single cell suspension. Spleens were process via grinding between frosted slide ends, dissociation via pipet, and passage through 70μM filter mesh to generate a single cell suspension. Peripheral blood was collected for analysis either through terminal cardiac puncture or tail vein. Flow cytometric identification of mature cell populations was performed using the following antibodies: APC-antiCD11b (BioLegend 101212), PE-Cy7-antiGr1 (BioLegend 108416), PERCP-Cy5.5-antiB220 (BioLegend 103236), and PE-antiCD3 (BioLegend 103206). Flow cytometric identification of hematopoietic stem and progenitor populations was performed using a combination of a lineage cocktail in addition to the following antibodies: APC-antiC-kit (BioLegend 105812), PERCP-CY5.5-antiSca1 (BioLegend 108124), AlexaFluor700-antiCD48 (BioLegend 103426), and PE-Cy7-antiCD150 (BioLegend 115914) with definition as annotated in text. Lineage cocktail consited of the following PE-conjugated antibodies, all from BioLegend: CD2 (100108), CD3 (100206), CD5 (100608), CD8 (100708), B220 (103208), Gr1 (108408), Ter119 (116208). Staining as performed for 30 minutes on ice in FACS buffer in the dark. Flow cytometry was performed using a BD FACSAria II or BD Fortessa. Peripheral blood counts were obtained using a Hemavet. Total nucleated cell numbers in the bone marrow and spleen were obtained using a Countess II with trypan blue.

### Competitive Bone Marrow Transplantation

Tester populations were maintained on the C57BL/Ka-CD45.2:Thy1.1. Competitor bone marrow cells were obtained from C57BL/Ka-CD45.1:Thy1.2 mice. Recipients were C57BL/Ka-CD45.2:Thy1.1X C57BL/Ka-CD45.1:Thy1.2 (F1) mice unless otherwise noted. Recipients received lethal irradiation with delivery of 550cGy twice with a minimum of 2hrs in between doses. Bone marrow was mixed in 1:1 ratio tester:competitor and delivered via retroorbital injection.

### BODIPY Staining

For C12 BODIPY staining following cell surface staining, samples were washed in HBSS with 1% FBS. Samples were then resuspended in 500uL HBSS with 0.5% fatty acid free BSA (Sigma) and incubated at 37C for fatty acid starvation. Stock C12 BODIPY (Cayman) was maintained at 1mg/ml. C12 BODIPY was then added to samples at 1:1000 dilution. Samples were then incubated for an additional 30 minutes at 37C. Samples were then washed 2 times in HBSS with 1% FBS, resuspended in 500uL of the same buffer, and immediately analyzed by flow cytometry.

For BODIPY 493/503 neutral lipid staining, samples were again cell surface stained as above and washed as above. BODIPY 493/503 (Cayman) was maintained as a 5mM stock. Cells were resuspended in 1mL HBSS with 1% FBS. BODIPY 493/503 was added at a 1:2000 dilution and incubated for 15 minutes at 37C. Samples were then washed twice and resuspended in 500uL of the same buffer, and immediately analyzed by flow cytometry using the BD Fortessa.

### Reactive Oxygen Species Measurement

Bone marrow cells were process and stained with cell surface markers as above. Following cell surface staining, samples were stained with either 5uM CellROX or 5uM CM-H2DCFDA for 30 minutes at 37C. Samples were then washed and analyzed immediately on BD Fortessa.

### Cell Cycle Analysis

For Ki67 staining, samples were collected as described in the “Hematopoietic Cell Isolation and Flow Cytometry” section. Following initial passage through 70μM filter mesh, cells were labeled with anti-CD117 microbeads (Miltenyi). Samples were then washed twice in FACS buffer and enriched using an AutoMACS NEO (Miltenyi). Samples were then stained with surface antibodies for hematopoietic stem and progenitors as above. Following staining, samples were washed. Samples were then fixed and permeablized according to manufacturer’s instructions using the transcription factor staining buffer set (eBioscience 00-5523-00). Samples were then stained in the dark at room temperature for 40 minutes using 10uL FITC-antiKi67 (Biolegend 652410). Samples were washed and then resuspended in 500uL permeabilization buffer with 5uL DAPI. Following a 20 minute incubation at room temperature in the dark, cells were washed twice in FACS buffer and analyzed on the BD Fortessa.

For EdU (5-ethynyl-2’-deoxyuridine) staining mice were injected with 1mg EDU (Cayman) and then maintained on water containing 0.3mg/mL EDU for 72hrs. Samples were prepared as above for flow cytometry through the fixation and permeabilization step. The click reaction was then performed using Alexa Fluor 488 Click-iT kit (Invitrogen C10641). DAPI staining was performed as above in Ki67 section followed by analysis on the BD Fortessa.

### Intracellular ATP Measurement

Bone marrow mononuclear cells were processed and stained to label LSK as above. LSK were then FACS sorted on the BD FACSAria II. LSK from each animal were then divided into technical triplicates of 10,000K cells each and ATP was measured with ATP Bioluminescence Assay Kit CLS II (Roche).

### Palmitate-Azide Click labeling

BM cells were stained with a Biotin lineage cocktail and washed before incubation with anti-Biotin Microbeads (Miltenyi Biotec). Following lineage depletion with the AutoMACS Neo(Miltenyi Biotec), cells were starved in IMDM + 1% fatty acid-free BSA for 30 min at 37C. BSA-palmitate or BSA-palmitate-azide were prepared by mixing equal volumes of 200 uM Palmitate or Palmitate Azide with 240 uM KOH and incubating in a 65C water bath for 15 min followed immediately by addition of IMDM + 1% fatty acid-free BSA. After starvation, cells were fed with either BSA-palmitate or BSA-palmitate-azide at a final concentration of 10 uM for 1 h. Cells were then stained for surface markers for 1h on ice and divided into equal groups for compartment-specific staining described previously.^22,32^ Briefly, cells were stained with one of 100 nM of BDP-DBCO for 15 min at RT, 100 uM of AlexaFluor488-DBCO for 30 min at 15C, or 50 nM of Cy5-DBCO for 15 min at 37C. Samples stained with Cy5-DBCO were then quenched with 100 nM of Cy7-N3 before 3 cycles of 15 min treatments at 37C with a high K+ wash buffer + 20 uM Valinomycin + 50 uM CCCP. All samples were washed at least 3 times before flow cytometric analysis.

All compounds were acquired from MedChemExpress

### Lipidomics

Samples were prepared for lipidomics and run in the Morisson lab as described previously.^9^ 10,000 CD48+LSK or CD48-LSK were used per run. Metabolomics data were then analyzed using the Omics Data Analyzer tool (<u>CRI / Omics Data Analyser · GitLab</u> developed by Zhiyu Zhao and described tin the cited reference. Samples were filtered to exclude all-zero metabolites and normalized using the Relative Log Expression (RLE) method. After normalization, missing values and zeros were imputed using a half of the minimum of the metabolites, and samples were log2-transformed. All the downstream analysis was performed using the preprocessed dataset. The Principal Component Analysis (PCA) was performed using all the preprocessed metabolites, and the top 2 components were selected for sample visualization. The heatmap was generated on the z-scores of all the FAs and carnitines. Differential expression tests were performed using Generalized Linear Mixed-Effects Models (GLMERs) on all the preprocessed metabolites. All analysis was performed using R4.5.0 with the stats, lme4, pheatmap, ggplot2, and ggforce packages.

### Statistical Analysis

Data are represented as mean +/-SD. N represents the number of biological replicates and is noted in individual figure legends. For samples comparing the means of two groups, two-tailed Student t-tests were performed. For comparisons of means between more than 2 groups, ANOVA was utilized. Differences attributable to mouse sex were analyzed, and since there were none, experiments represent samples pooled with near balanced numbers of each sex. Experiments were performed at least twice and data were pooled. Mouse numbers for each experiment were not based on power calculations, but were based on lab experience dependent on the assay performed.

