## Supplemental Figure Legends for "ACC1-Dependent De Novo Lipogenesis Sustains Hematopoietic Stem Cell Quiescence and Self-Renewal"

**Supplemental Figure S1. Confirmation of ACC1 deletion following poly(I:C) injection and hematopoietic progenitor analysis.**

(A) Western blot analysis of cKit+ progenitors from WT and ACC1 KO mice 4 weeks after poly (I:C) injection.

(B) qPCR analysis of *Acaca* gene expression in FACS sorted LSK progenitor in WT and ACC1 KO mice 4 weeks after poly (I:C) injection.

(C) Frequency of common myeloid progenitor (CMP;Lineage^-^cKit^+^CD34^+^FcgR^-^), granulocyte-monocyte progenitor (GMP;Lineage^-^cKit^+^CD34^+^FcgR^+^), and megakaryocyte-erythrocyte progenitor (MEP;Lineage^-^cKit^+^CD34^-^FcgR^-^), populations in WT and KO mice.

**Supplemental Figure S2. Vav1-Cre-mediated deletion of ACC1 results in fetal demise and disrupted hematopoiesis.**

(A) e15.5 ACC KO fetuses are pale compared to wildtype littermates.

(B) Comparison of fetal liver cellularity in ACC WT, heterozygous (Het) and KO fetal liver at e15.5.

(C) Comparison of myeloid cells (CD11b+) in fetal livers at e15.5.

(D) Comparison of B cells (B220+) in fetal livers at e15.5.

(E) Comparison of LSK cells in fetal livers at e15.5.

(F) Comparison of HPC (CD48+ LSK) cells in fetal livers at e15.5.

(G) Comparison of MPP (CD48-CD150-LSK) cells in fetal livers at e15.5.

(H) Comparison of LT-HSC (CD48-CD150+LSK) in fetal livers at e15.5.

In the above figure data are derived from at least 4 individual mice per group with values from individual mice shown. Graphs display mean + S.D. Anova was used for comparison of means. P values *<0.05, **<0.01, ***<0.001, ****<0.0001.
