## Supplementary figures and images for "ACC1-Dependent De Novo Lipogenesis Sustains Hematopoietic Stem Cell Quiescence and Self-Renewal"

### supplemental figures 1 and 2

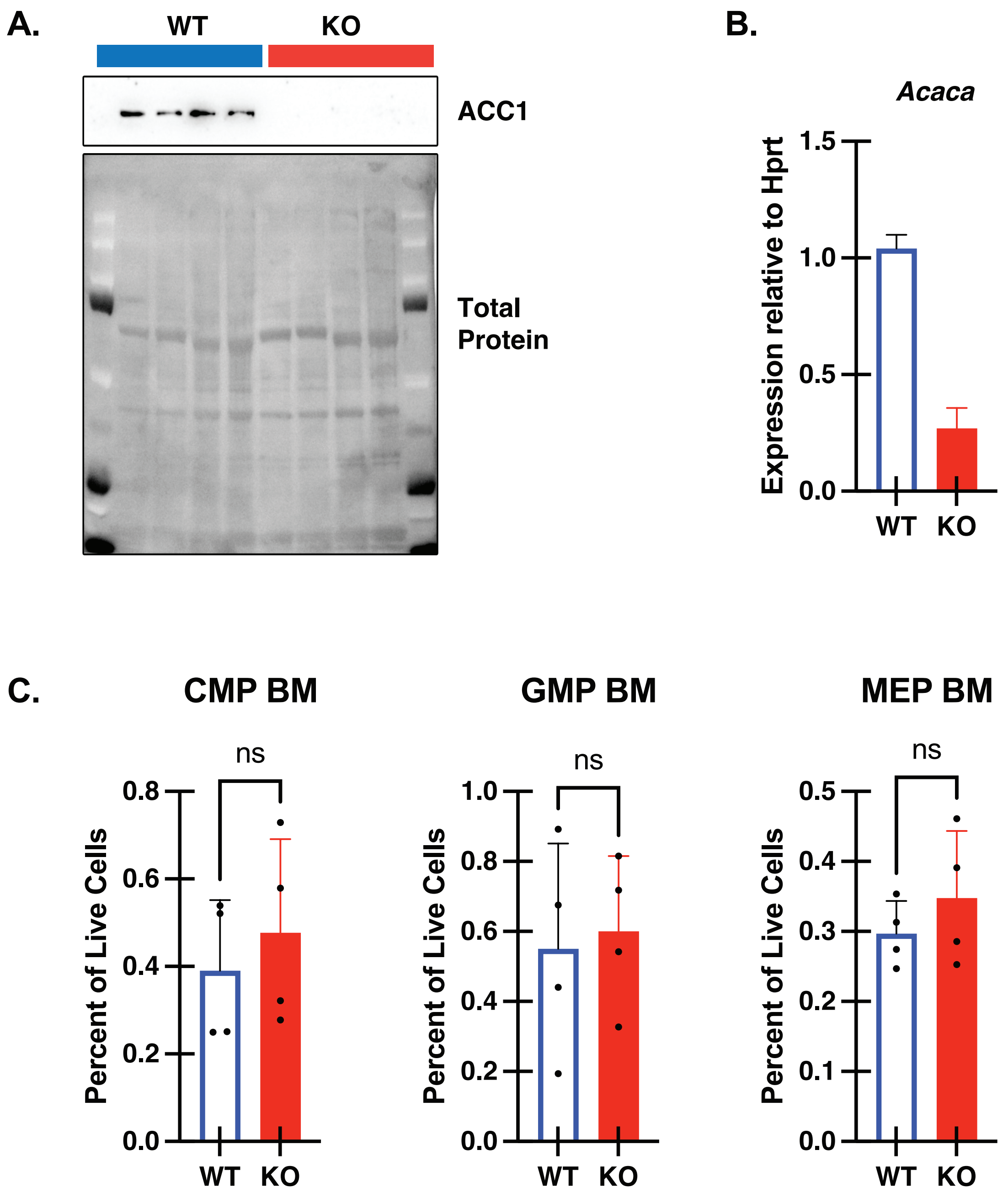


Supplemental Figure 1

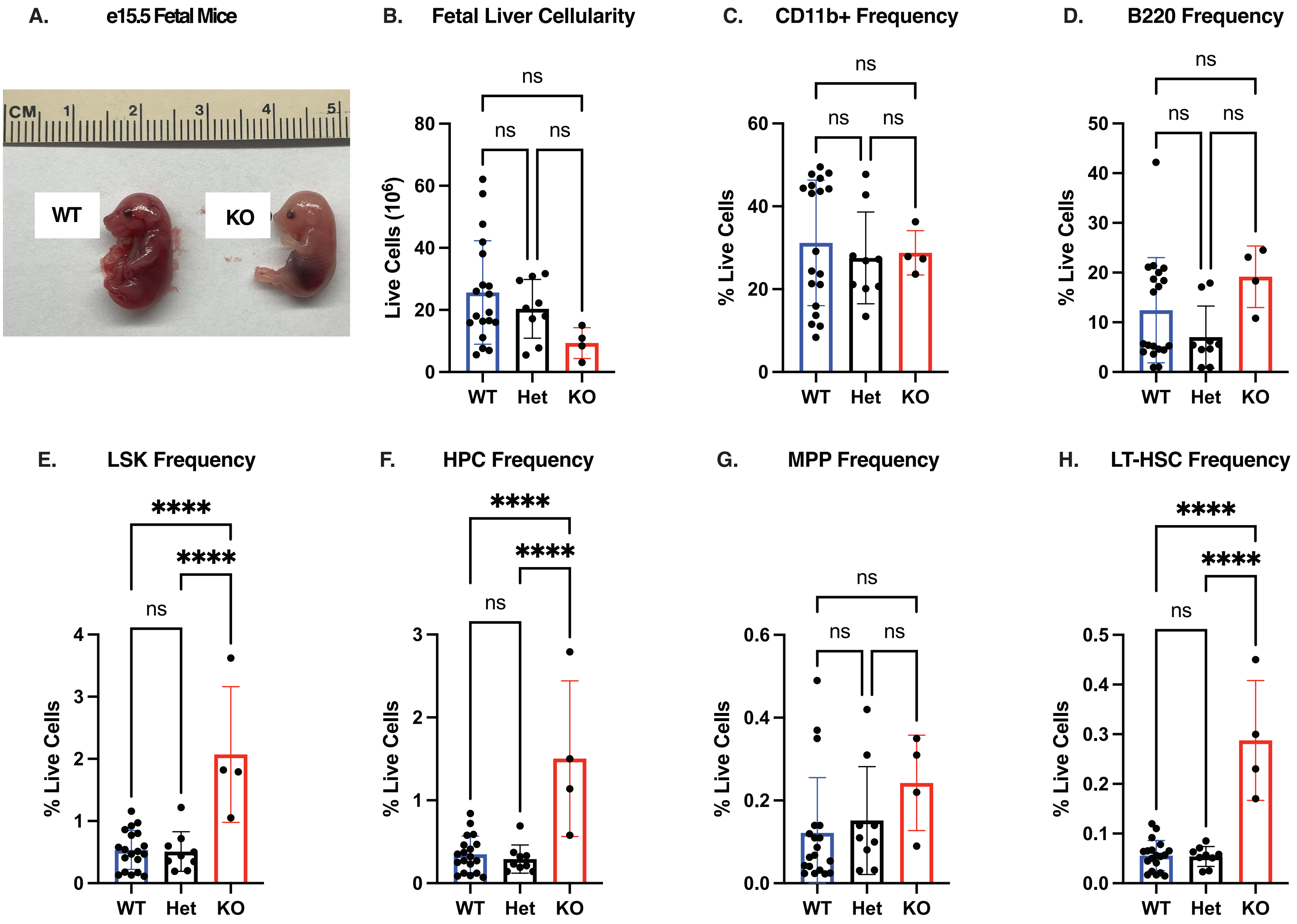


Supplemental Figure 2
